# 3D reconstruction of spatial transcriptomics with spatial pattern enhanced graph convolutional neural network

**DOI:** 10.64898/2026.01.13.699328

**Authors:** Chen Tang, Yuansheng Zhou, Xue Xiao, Lei Dong, Lei Yu, Qiwei Li, Guanghua Xiao, Lin Xu

## Abstract

Spatially resolved transcriptomics (SRT) is a promising new technology that enables simultaneous analysis of gene expression and spatial information for biomedical research. However, the existing statistical and deep learning algorithms used for analyzing SRT data rely solely on two-dimensional (2D) spatial coordinates, which limits their ability to accurately identify spatial domains, spatially variable genes, cell-to-cell communications, and developmental trajectories in a three-dimensional (3D) spatial manner. To address these limitations, we introduced Spa3D, which utilized the anti-leakage Fourier transform and graph convolutional neural network model to reconstruct 3D-based spatial structures from multiple 2D SRT slices. We demonstrate that Spa3D is appliable to analyze data from various SRT technology platforms and outperforms state-of-art methods by: (I) improving spatial domain identification through 3D reconstruction, (II) elucidating cell-cell communication landscape in the 3D cellular organization, (III) modeling of organ-level tempo-spatial development patterns in a 3D fashion, and (IV) annotating 3D spatial trajectory that are not captured by 2D spatial coordinates.

**Key points:**

- Most existing spatial omics analysis methods rely on 2D data, limiting their ability to capture full spatial and developmental tissue complexity
- Spa3D incorporates physical z-axis distances, enabling accurate 3D modeling even when adjacent slices vary in tissue structure and composition
- Spa3D reconstructs true 3D spatial structures from 2D SRT slices using graph convolutional networks and anti-leakage Fourier transforms
- Spa3D enhances spatial domain detection, revealing detailed cell-cell communication and organ-level development patterns across multiple spatial omics platforms
- Spa3D enables novel biological discoveries by revealing spatial features and trajectories not detectable using traditional 2D transcriptomic analysis approaches

## INTRODUCTION

Single-cell transcriptomics (scRNA-seq) has revolutionized biomedical research, enabling exploration of gene expression at the individual cell level. However, most scRNA-seq protocols destroy spatial context information that offers insights into tissue organization, cell–cell communication, and microenvironment^1,2^. To address this, spatially resolved transcriptomics (SRT) technologies^2–7^ integrate gene expression with two-dimensional (2D) spatial coordinates. This approach for biomedical studies^1,2^ creates an urgent need for computational methods that incorporate both 2D and three-dimensional (3D) spatial information into SRT data analysis to understand biological processes in tissue morphology.

Why do we need 3D information for spatial transcriptomics? Most biological processes occur in three-dimensional tissue structures that enable cell-cell interactions, communication, and specialized microenvironments. A single 2D slice cannot capture these complex 3D relationships. For example, a liver cross-section may intersect tubular central veins at varying angles, producing different lumen shapes. Thus, understanding spatial dynamics and interactions in 3D is essential for fully interpreting biological function.

Advances in statistics and deep learning have offered insights into SRT data—identifying spatial domains^8–10^, cell-cell communication^11–16^, and developmental trajectories^17–22^. However, these computational algorithms^8–23^ rely on 2D coordinates, preventing them from leveraging true 3D organization. For instance, SpaGCN^8^ combined gene expression with image data for spatial clustering but only used 2D coordinates. Additionally, methods such as CellPhoneDB^11^, CellChat^12^, and NicheNet^13^ were designed for single-cell RNA-seq. Newer tools for SRT (e.g. SpaTalk^14^, COMMOT^15^, and NCEM^16^) also limit themselves to 2D spatial coordinates without incorporating 3D interaction information. Similarly, existing pseudotime and trajectory analysis methods either omit spatial information^18–22^, or only leverage 2D coordinates^17^. These limitations motivate the need for algorithms that exploit 3D data to define spatial domains, cell–cell communication networks, and developmental trajectories, thereby maximizing SRT’s potential.

PASTE^24^ is a leading method for aligning multiple 2D SRT slices in 3D. It offers two modes: pairwise slice alignment, which stacks slices without reconstructing true 3D gene expression, and center slice integration, which merges data into a single 2D “center” slice—sacrificing 3D resolution. PASTE has three major limitations. First, its “center slice” provides only a 2D overview rather than a genuine 3D reconstruction, so it cannot deliver slice-specific 3D clustering or trajectory information. Second, PASTE ignores real z-axis distances between adjacent slices, preventing accurate 3D layout reconstruction. This issue also affects other integration methods such as DeepST^25^ and BASS^26^, which use deep learning or Bayesian models for segmentation instead of explicit 3D reconstruction. Third, PASTE assumes that adjacent 2D slices are highly similar, which often fails on real datasets where neighboring sections differ.

To overcome these challenges, we developed Spa3D, a graph-convolutional neural network (GCN) for 3D reconstruction from multiple 2D SRT slices that explicitly incorporates true z-axis distances, enabling more accurate 3D reconstructions than PASTE. By using actual x-, y-, and z-axis coordinates, Spa3D constructs a genuine 3D structure rather than relying on a 2D “center slice.” We applied Spa3D to various SRT platforms (e.g., 10x Genomics Visium, ST, Stereo-seq, and MERFISH) and found that it can (I) improve spatial domain identification, (II) reveal detailed 3D cell–cell communication networks, (III) map organ-level temporal and spatial development in 3D, and (IV) identify 3D spatial trajectories that are invisible in 2D. Overall, Spa3D provides a fuller 3D picture of cellular organization and development, with significant implications for biomedical research and clinical applications.

## RESULTS

### Overview of Spa3D

**Figure 1A** illustrates the overall workflow of Spa3D, which integrates consecutive multi-slice 2D spatial transcriptomics (SRT) datasets for 3D reconstruction and downstream analyses. Each 2D SRT dataset is first processed using either the Hilbert transform or the anti-leakage transform to extract coherent spatial signals and suppress noise, thereby enhancing biologically meaningful spatial features within individual slices prior to alignment or graph learning. The enhanced 2D omics datasets are then aligned to correct inter-slice deviations and generate aligned spatial coordinates (x’,y’). After alignment, z-coordinates are assigned to each slice based on their physical or sequential order. The aligned (x’,y’) coordinates and assigned z-positions are combined to construct a 3D adjacency matrix that encodes spatial relationships among all points across slices and modalities, capturing both local 2D neighborhood structure and inter-slice proximity. This adjacency matrix, together with node features from all enhanced omics, is subsequently input into a Graph Convolutional Network (GCN). The GCN learns low-dimensional embeddings, which are further refined through a Deep Embedded Clustering (DEC)-based optimization using soft cluster assignments. In this way, the GCN model integrates molecular and spatial information to produce spatially coherent tissue domains. Collectively, Spa3D outputs 3D spatial domain maps, aligned and denoised multi-omics feature matrices, and learned embeddings, which support diverse downstream analyses such as spatial domain identification, pseudotime inference, cell-cell communication and 3D spatial trajectory **(Figure 1B**).

**Figure 1.**
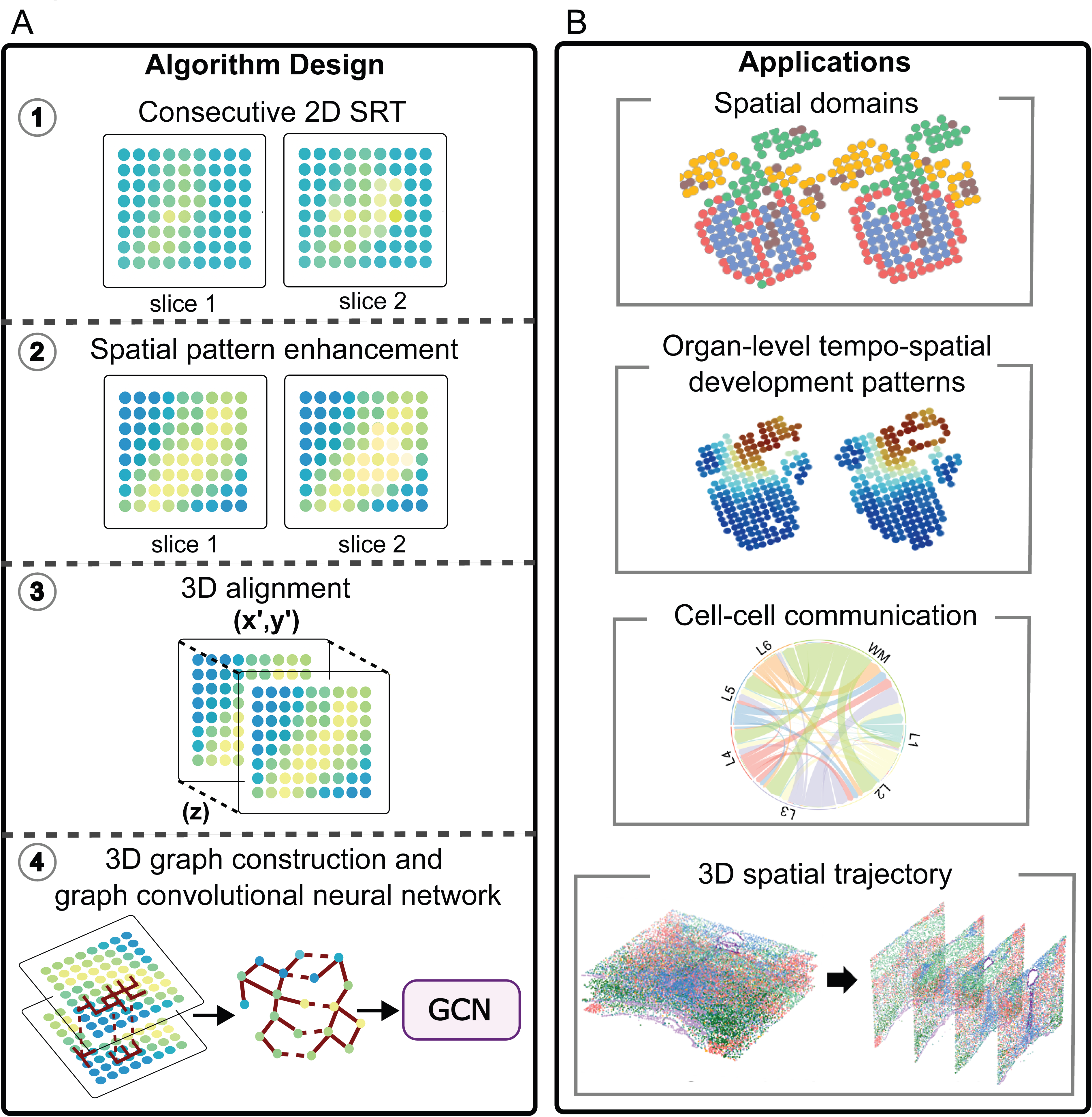
Overview of the Spa3D workflow and its applications. **(A)** The schematic of Spa3D workflow which comprises several key steps. First, multiple consecutive 2D spatial transcriptomics (SRT) slices are collected as input. Second, each slice undergoes spatial pattern enhancement using the anti-leakage Fourier transform (ALFT) algorithm to extract coherent spatial features and suppress noise. Third, the spatial coordinates are aligned with the techniques implemented in PASTE if the aligned coordinates are not available. After assigning z-coordinates based on slice order, a 3D graph network is constructed to capture both intra- and inter-slice spatial relationships. The graph and molecular features are then input into a graph convolutional network (GCN) trained with a DEC-inspired clustering strategy to generate integrated embeddings and spatially coherent tissue domains. (**B**) The four major applications of Spa3D: identification of spatial domains, elucidation of organ-level tempo-spatial developmental patterns in three dimensions, mapping of cellular communication landscapes, and discovery of 3D spatial trajectories.

### Human dorsolateral prefrontal cortex dataset by 10X Genomics Visium platform

To compare Spa3D with PASTE, we applied Spa3D to the human dorsolateral prefrontal cortex (DLPFC) dataset^27^, previously used to showcase the capability of PASTE in 3D spatial domain identification^24^. This 10x Genomics Visium dataset uses curated annotations from the original study^27^ as ground truth.

First, to assess spatial-domain identification, we compared Spa3D and PASTE using the adjusted Rand Index (ARI). A higher ARI indicates closer agreement with the curated annotations^27^. In the original PASTE publication^24^, PASTE established stacked 3D alignment and integrated multiple SRT slices of DLPFC samples into a single “center slice” using optimal transport formulation. In contrast, Spa3D characterizes the detailed 3D cellular organization in each individual SRT slice without the need to establish a single 2D-based “center slice”. Across all DLPFC slices in the 3D spatial reconstruction, Spa3D achieved an ARI score of 0.62 for the whole 3D spatial structure as depicted in **Figure 2A**. Because the Spa3D directly presented the 3D structure while the PASTE method only provided a summarized 2D “center slice” to represent all the slices, for a fair comparison purpose, we compared the 2D “center slice” (ARI = 0.53, **Figure 2B**) with the corresponding 2D slice from Spa3D reconstruction (ARI = 0.63, **Figure 2B**), instead of comparing with the whole Spa3D-defined 3D reconstruction in **Figure 2A**. It is important to note that all of the Spa3D-reconstructed individual slices (ARI from 0.58 to 0.63) (**Supplementary Figure 1**) outperformed the PASTE-defined “center slice” in clustering accuracy, not just the one showcased in **Figure 2B**.

**Figure 2.**
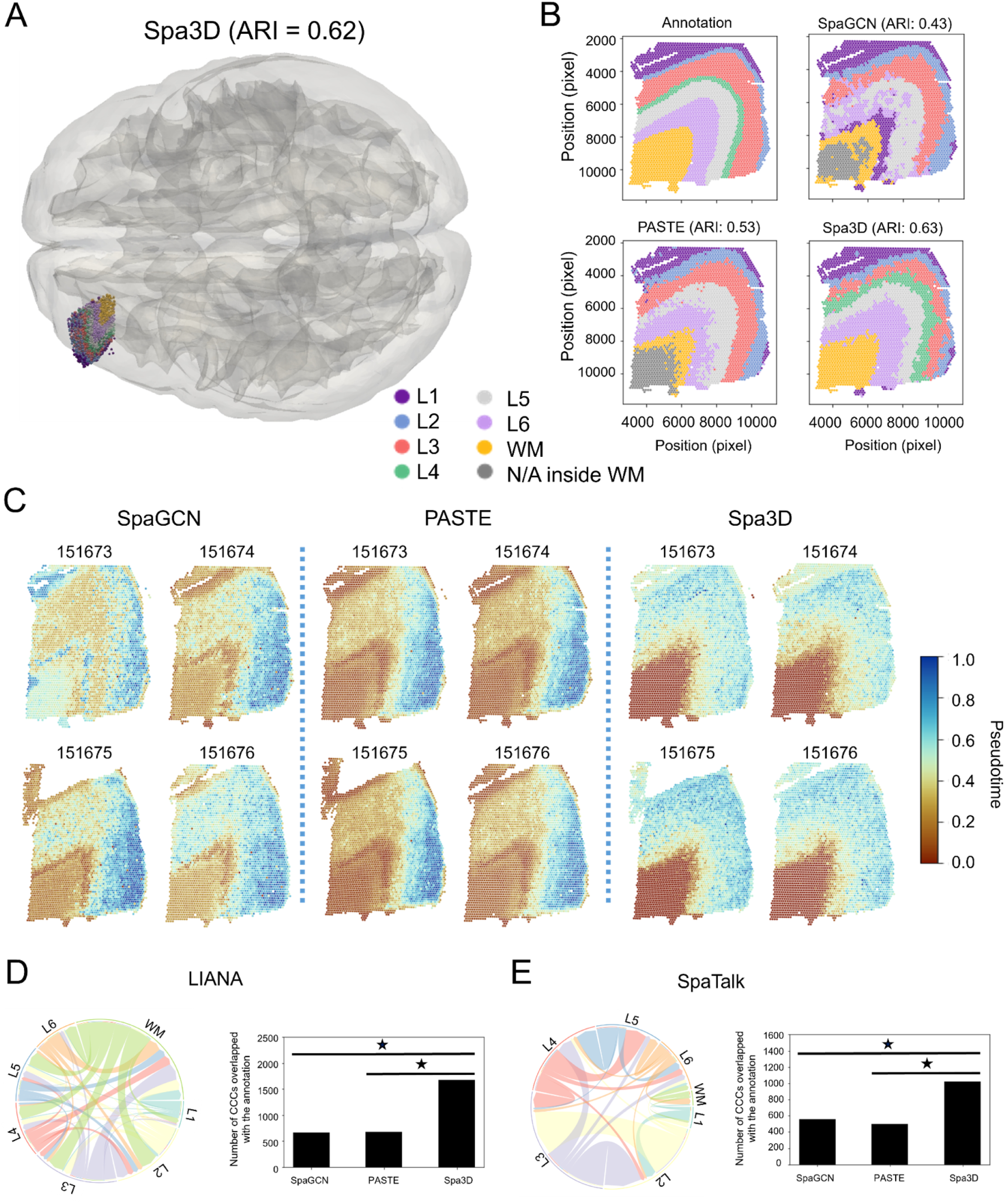
3D characterization of the human dorsolateral prefrontal cortex dataset. **(A)** Reconstruction of the 3D cellular organization by Spa3D, with incorporation into a human brain structure for descriptive purposes. The IDs of the 2D SRT slices used in this analysis are 151673, 151674, 151675, and 151676, respectively. The third dimension was based on real spatial coordinates. ARI was calculated based on 3D instead of 2D information. **(B)** Comparisons of spatial domains among the annotation, SpaGCN, PASTE, and Spa3D. **(C)** Comparisons on the pseudotime estimated from SpaGCN, PASTE, and Spa3D results. **(D-E)** Cell-cell communication (CCC) analysis based on LIANA (D) and SpaTalk (E). PASTE- and Spa3D-defined clustering results were passed to LIANA and SpaTalk as inputs. The chord diagram (left) illustrated the cell-cell communication frequencies among clusters, while the bar plot (right) showed the number of ligand-receptor pairs overlapped with the annotation. Statistical significances were determined by a two-tailed Fisher exact test. *P < 0.05. The true positive rate (TPR) and false positive rate (FPR) for ligand-receptor pairs (FPLRPs) for Panels E and F are shown in Supplementary Figure 4.

To understand whether utilizing 3D spatial reconstruction via Spa3D will improve 2D-based spatial domain analysis, we compared its performance with that of SpaGCN, the state-of-the-art spatial domain segmentation algorithm for 2D SRT data. While Spa3D-defined spatial domains and curated annotations shared the same layer or domain structures, SpaGCN-defined spatial domains missed a true layer (green in the curated annotation figure) and introduced an artificial layer (grey in the SpaGCN figure) that did not match with the curated annotations (ARI = 0.43, **Figure 2B**). Additionally, we observed several inconsistencies between PASTE-defined spatial domains and curated annotations, including a missing layer (green in the annotation figure) and a wrongly annotated layer (grey in the PASTE figure) that did not exist in the annotation figure, both of which negatively impacted its clustering accuracy (**Figure 2B**). These results show that Spa3D outperforms other methods in identifying spatial domains and that using 3D reconstruction improves domain segmentation. Besides SpaGCN and PASTE, we also compared Spa3D to another two competing methods STAGATE and GraphST in **Supplementary Figure 2** and showed that Spa3D achieves the highest ARI.

To examine whether the identified domains from Spa3D are biologically meaningful, we extracted <u>s</u>patial <u>d</u>omain <u>a</u>ssociated <u>s</u>patially <u>v</u>ariable <u>g</u>enes (sdaSVGs), which were defined previously^8^ by differential expression analysis among different spatial domains. To achieve a reliable and robust conclusion, we applied two most commonly used approaches, edgeR^28^ and Seurat^29^ packages, to present the results of differential expression analysis on the seven layers (L1 – L6 and WM) of the prefrontal cortex (**Supplementary Figure 3).** We compared Spa3D with PASTE in identifying sdaSVGs of each of the seven layers in **Figure 2A** and identified sdaSVGs overlap with the known dorsolateral prefrontal cortex gene set from the Allen Mouse Brain Atlas^30^. Our analysis revealed that, in the majority of comparisons, Spa3D-defined sdaSVGs are more enriched of known dorsolateral prefrontal cortex genes than PASTE-defined sdaSVGs, as indicated by both edgeR (**Supplementary Figure 3A**) and Seurat (**Supplementary Figure 3B**) approaches. This consistency between two distinct differential expression analysis methods suggests that the spatial domains and domain-specific expressed genes identified by Spa3D are more biologically meaningful and consistent with existing knowledge than PASTE. Overall, our results demonstrate the consistent superiority of Spa3D in detecting spatial domains and sdaSVGs.

Second, we evaluated whether Spa3D-derived pseudotime could be more accurate than pseudotime inferred based on PASTE and SpaGCN (**Figure 2C**). Across all four SRT slices, Spa3D-derived pseudo-temporal structure corresponds to the well-established “inside-out” layer pattern of corticogenesis^31,32^: new neurons are born in the ventricular zone, migrate along the radial glia fibers in a vertical fashion towards the marginal zone. Therefore, Spa3D’s pseudotime accurately reflects the dorsolateral prefrontal cortex’s layered organization, whereas PASTE and SpaGCN fail to capture the inside−out sequence of cortical layer development. This highlights how Spa3D’s 3D reconstruction improves pseudotime inference, offering deeper insights into the molecular mechanisms of development and disease.

Third, we investigated whether Spa3D could accurately identify ligand-receptor signaling pathways involved in cell-cell communication, in comparison with PASTE and SpaGCN (**Figures 2D and 2E**). To robustly define cell-cell communication, we employed two distinct frameworks, LIANA^33^ and SpaTalk^14^. The LIANA package comprises several most frequently used cell-cell communication analysis algorithms that were originally designed for scRNAseq data without considering spatial information, such as CellChat^12^ and CellPhoneDB^11^. In contrast, SpaTalk is a new algorithm designed for elucidating spatially resolved cell-cell communications using SRT data. We applied both LIANA and SpaTalk to define the cell-cell communication landscape based on spatial clusters identified by the curated annotations from the original study^27^, Spa3D, PASTE, and SpaGCN. To assess how well each approach aligned with the curated annotations on cell-cell communications, we calculated the overlapping cell-cell communications between the annotation and each of the three methods examined here (Spa3D, PASTE, and SpaGCN). Specifically, SpaGCN identified 1,976 cell-cell communications in total, with 661 overlapping with annotation (Jaccard index = 0.197); PASTE identified 1,953 cell-cell communications in total, with 678 overlapping with annotation (Jaccard index = 0.204); Spa3D identified 2,075 cell-cell communications in total, with 1,679 overlapping with annotation (Jaccard index = 0.689) (**Figure 2D**). Interestingly, we found that, based on the LIANA framework, Spa3D identified 1679 ligand-receptor interacting pairs that overlapped with the annotation, which was more than twice the number of overlaps between the annotation and PASTE or SpaGCN (**Figure 2D**). A similar trend was observed using the SpaTalk framework (**Figure 2E**). SpaGCN identified 2,277 cell-cell communications in total, with 558 overlapping with annotation (Jaccard index = 0.164); PASTE identified 1,826 cell-cell communications in total, with 501 overlapping with annotation (Jaccard index = 0.167); Spa3D identified 2,303 cell-cell communications in total, with 1,024 overlapping with annotation (Jaccard index = 0.346). These findings indicate that the 3D spatial construction via the Spa3D algorithm captures the nature of cell-cell communication more accurately than the other methods. **Supplementary Figure 4** provides a false positive rate (FPR) for ligand-receptor pairs and confirms the same trend. These findings demonstrate that Spa3D can identify cell-cell communication networks and providing important insights into the cellular interaction mechanisms underlying spatially resolved biological processes.

### Human embryonic heart dataset by barcoded solid-phase RNA capture for Spatial Transcriptomics (ST) profiling platform

To gain a comprehensive understanding of how organs develop and how diseases progress in 3D spatial biology, it is necessary to extract comprehensive 3D knowledge at the organ level^1,34,35^. To demonstrate the capability of Spa3D in delineating 3D spatial domains at the whole organ level, we applied it to the human embryonic heart dataset^36^ at 6.5 post-conception weeks (PCW) (4 sections) generated by barcoded solid-phase RNA capture for spatial transcriptomics profiling platform^37,38^. Our findings, as depicted in **Figure 3A**, reveal that the 3D cellular organization defined by Spa3D accurately reflects the heart structure as annotated by Asp et al. (2019)^37^ (ARI = 0.48 for the 3D spatial structure in **Figure 3A**; ARI scores range from 0.44 to 0.57 for each individual slice calculated by Spa3D), thus confirming the capability of Spa3D to accurately capture the spatial organization at the organ level.

**Figure 3.**
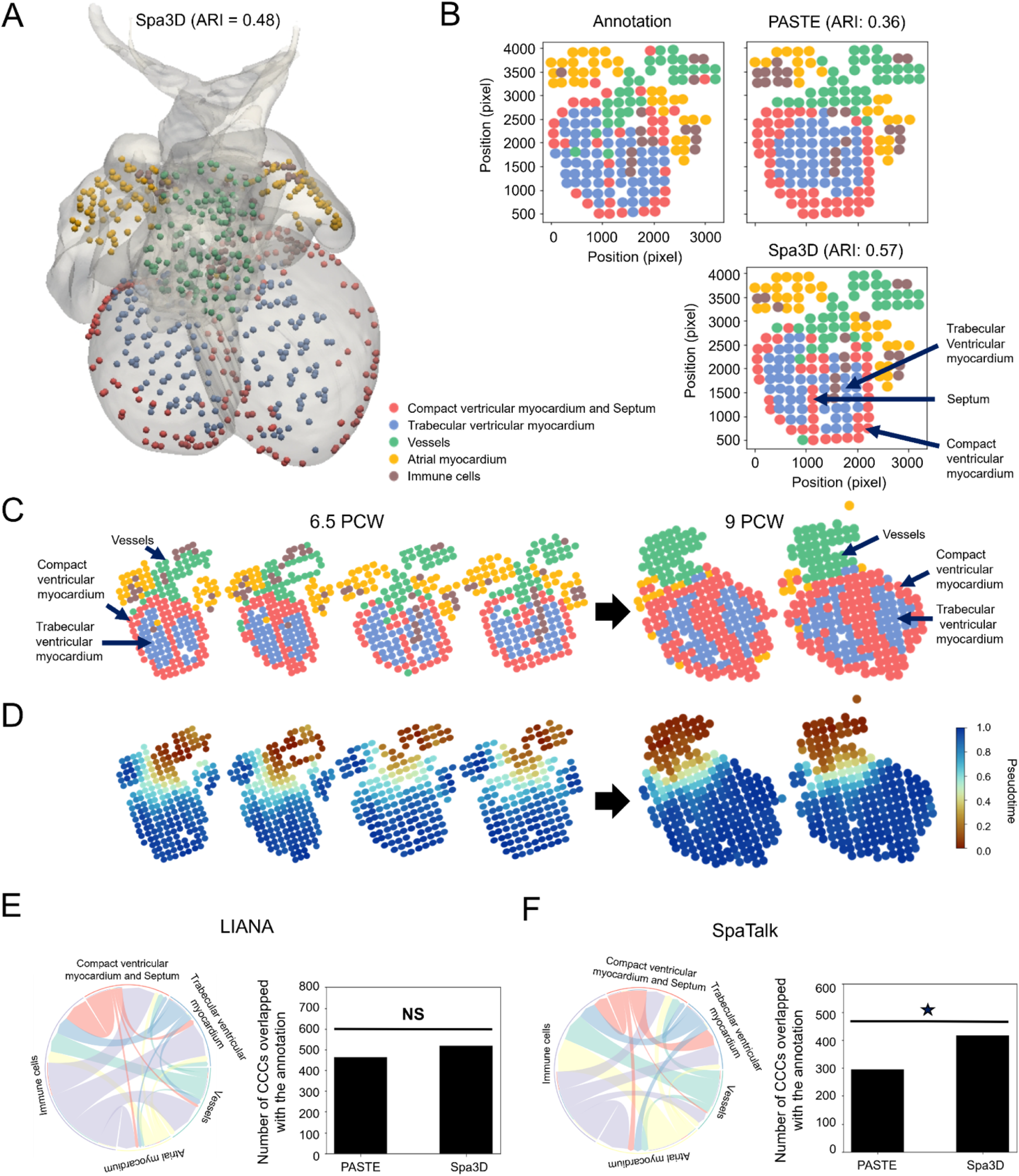
3D characterization of tempo-spatial development patterns in the human embryonic heart dataset. **(A)** Reconstruction of the 3D cellular organization by Spa3D, with incorporation into a human heart structure for descriptive purposes. The IDs of the four slices are 9, 10, 11, and 12, respectively. **(B)** Comparisons of spatial domains among the annotation, PASTE, and Spa3D. **(C)** Spa3D reconstructed cellular organization from 6.5 PCW to 9 PCW. **(D)** Spa3D-defined organ-level tempo-spatial development patterns from 6.5 PCW to 9 PCW. The six slices at 6.5 PCW and 9 PCW are processed together to present a consistent pseudotime. **(E-F)** Cell-cell communication (CCC) analysis based on LIANA (E) and SpaTalk (F). PASTE- and Spa3D-defined clustering results were passed to LIANA and SpaTalk as inputs. The chord diagram (left) illustrated the cell-cell communication frequencies among clusters, while the bar plot (right) showed the number of ligand-receptor pairs overlapped with the annotation from single-cell mapping. Statistical significances were determined by a two-tailed Fisher exact test. NS for non-significant. *P < 0.05. The true positive rate (TPR) and false positive rate (FPR) for ligand-receptor pairs (FPLRPs) for Panels E and F are shown in Supplementary Figure 8.

To ensure a fair comparison between the 3D cellular organization defined by Spa3D and PASTE-identified “center slice” on spatial domain segmentation, we first showcased a comparison between the PASTE-integrated 2D “center slice” (ARI = 0.36, **Figure 3B**) and the corresponding slice (ARI = 0.57, **Figure 3B**) from Spa3D reconstruction. Furthermore, we compared all individual slices reconstructed by Spa3D (ARI from 0.44 to 0.57) (**Supplementary Figure 5**) with PASTE’s “center slice” and found that Spa3D consistently achieved higher ARI scores than PASTE, not just the one showcased in **Figure 3B**. In addition, it is important to note that PASTE failed to show the heart septum between the left and right ventricles, a crucial structural element in the heart (**Figure 3B**). In contrast, Spa3D was able to display the septum structure between the left and right ventricles, consistent with our knowledge of heart anatomy and physiology^39–42^. Beyond comparison with the PASTE integration, we also evaluated Spa3D against two other methods, STAGATE and GraphST. **Supplementary Figure 6** demonstrates that Spa3D attained a higher ARI.

To explore domain-specific differentially expressed genes, we applied edgeR and Seurat packages on differential expression analysis to detect sdaSVGs based on Spa3D- and PASTE-defined spatial domains. As shown in **Supplementary Figure 7**, in most of the spatial domains, the sdaSVGs detected by Spa3D showed higher overlap with known heart-specific gene set than the ones defined by PASTE, confirming that Spa3D-defined sdaSVGs align better with known biological knowledge.

We next assessed the ability of Spa3D to elucidate the organ-level tempo-spatial development patterns. We aimed to comprehend the spatial and temporal developmental dynamics of the human embryonic heart. To achieve this, we compared Spa3D-defined 3D reconstructions at 6.5 post-conception weeks (PCW) (4 sections with IDs from 9 to 12) and 9 PCW (2 sections with IDs from 16 to 17). **Figure 3C** illustrates the Spa3D-defined spatial domain dynamics from 6.5 to 9 PCW. We observed significant changes in spatial domains during the heart development process, including the increase in the size of the spatial domain representing vessels and outflow tracts (green color) and the spatial domain for the compact ventricular myocardium and septum (red color). These findings are in line with current knowledge of human heart anatomy and physiology from 6.5 to 9 PCW^39,42–44^.

To continue the investigation of the temporal dynamics of organ-level development patterns, we performed the pseudotime inference analysis by DPT method^21^ at 6.5 and 9 PCW to infer the pseudo-spatial trajectories at these two time points based on Spa3D results (**Figure 3D**). Interestingly, while two distinct spatial domains (**red and blue in Figure 3C**) were observed in the ventricular myocardium, these two domains were indistinguishable with similar pseudotime scores in **Figure 3D**. We resolved this seeming contradiction through a deep exploration of the available anatomy and physiology evidence. Specifically, both spatial locations and molecular makers of these two domains in **Figure 3C** (e.g., IGFBP3, LDHA, PGK1, and ENO1 for red domain, and MYL2, MYH7, FHL2, and PLN for blue domain) aligned well with known compact (red) and trabecular (blue) ventricular myocardium structures in human hearts^36,45–47^. This result confirms that the Spa3D-defined spatial domains in **Figure 3C** are consistent with known cell types. In contrast, **Figure 3D** indicates that these two distinct cell types share similar pseudotime scores and temporal trajectory patterns. These results agree with experimental evidence that the formation and specification of compact and trabecular ventricular myocardium both occur during the same time period (weeks 5-8) of human embryonic life, as part of normal cardiac morphogenesis^45–49^. Collectively, these results support that 3D reconstruction via Spa3D can assist the spatial segmentation and pseudotime inference analysis to achieve good agreement with existing knowledge.

Finally, we investigated the capability of Spa3D to accurately identify ligand-receptor signaling pathways involved in cell-cell communication in comparison with PASTE. To evaluate their performance, we calculated the percentage of overlapping ligand-receptor pairs between the annotated from Asp et al. (2019)^37^ and Spa3D or PASTE results (**Figures 3E and 3F**). Under LIANA framework (**Figure 3E**), PASTE identified 702 cell-cell communications in total, with 464 overlapping with annotation (Jaccard index = 0.437); Spa3D identified 771 cell-cell communications in total, with 519 overlapping with annotation (Jaccard index = 0.483). Under SpaTalk framework (**Figure 3F**), PASTE identified 674 cell-cell communications in total, with 295 overlapping with annotation (Jaccard index = 0.209); Spa3D identified 800 cell-cell communications in total, with 415 overlapping with annotation (Jaccard index = 0.293). Therefore, under both LIANA and SpaTalk framework, Spa3D identified more ligand-receptor interacting pairs that overlapped with the annotation than PASTE. A statistically significant difference was observed under the SpaTalk framework (**Figure 3F**), while LIANA showed the same trend but was not statistically significant. LIANA’s lack of spatial information likely explains its nonsignificant results and lower accuracy versus SpaTalk for spatial transcriptomics. **Supplementary Figure 8** shows false-positive rates (FPR) for ligand-receptor pairs, confirming this trend. Thus, Spa3D effectively enhances cell-cell communication analysis via 3D reconstruction.

### High resolution developing *Drosophila* embryos dataset by Stereo-seq platform

To assess the performance of Spa3D on high-resolution spatial transcriptomic data, we analyzed a dataset^50^ derived from developing *Drosophila* embryos using Stereo-seq, which captured the expression of 13,668 genes at near-single-cell resolution. The dataset includes 16 consecutive slices sampled at uniform intervals covering the entire embryo during stage E14–16 (**Figure 4A**). For 3D reconstruction, we selected four representative slices (slices 9–12). Spa3D well reconstructed the 3D architecture of these slices (**Figure 4B**, ARI = 0.29). Using slice 11 as the center slice, we found that Spa3D (ARI = 0.38) outperformed PASTE (ARI = 0.27) in domain segmentation (**Figure 4C–E**). In particular, Spa3D more accurately identified tracheal and CNS cells compared with PASTE (**Fig. 4D–E**).

**Figure 4.**
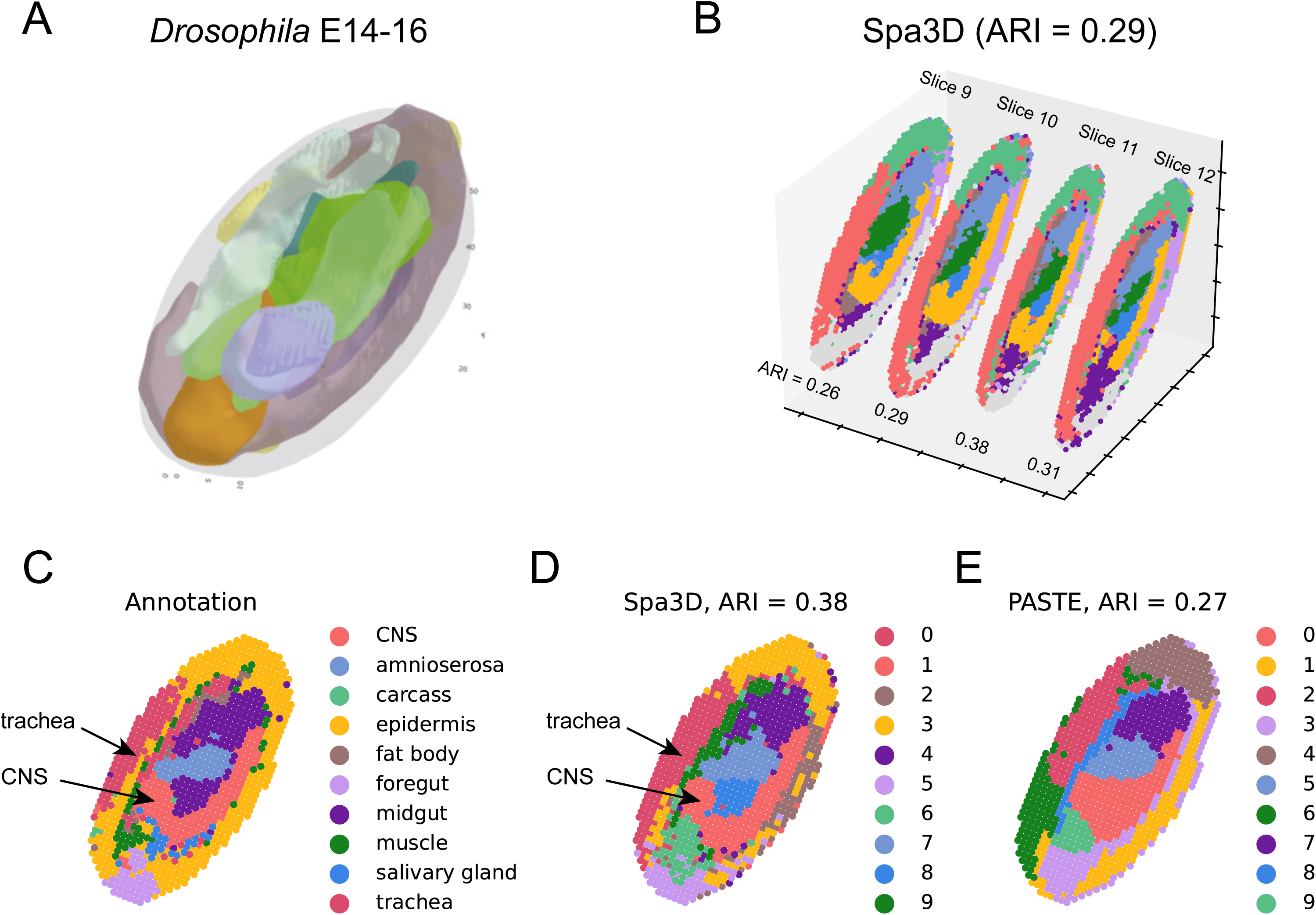
3D characterization of developing Drosophila embryos dataset. **(A)** The schematic of 3D Drosophila embryos structure at stage E14-16. (**B**) Reconstruction of the 3D structure of slices 9-12 by Spa3D. (**C**) Annotation of cells in slice 11. (**D**) Doman segmentation of slice 11 by Spa3D. (**E**) Doman segmentation of slice 11 by generating center slice with PASTE.

### 3D spatial trajectory in the mouse hypothalamus dataset by MERFISH platform

To demonstrate the versatility of Spa3D in analyzing SRT data from various platforms, we studied a dataset^3^ from the mouse hypothalamic preoptic region generated using the multiplexed error-robust fluorescence in situ hybridization (MERFISH) approach. This dataset is distinct from the 10X Genomics Visium, ST and Stereo-Seq platforms presented in **Figure 2-4**, respectively, and represents an imaging-based singe-cell resolution SRT approach.

First, we conducted a comparative analysis of Spa3D and PASTE on this mouse hypothalamus dataset to assess their ability to accurately identify spatial domains. Our results showed that Spa3D not only accurately reflected the anatomical structure of the hypothalamus as annotated by Moffitt et al. (2018)^3^ (ARI = 0.50 for the whole 3D reconstruction, ranging from 0.49 to 0.53 for each individual SRT slice calculated by Spa3D, **Figure 5A**, with a planar layout in **Supplementary Figure 9**), but also outperformed PASTE-defined central slice in clustering accuracy (ARI = 0.04, **Supplementary Figure 10**).

**Figure 5.**
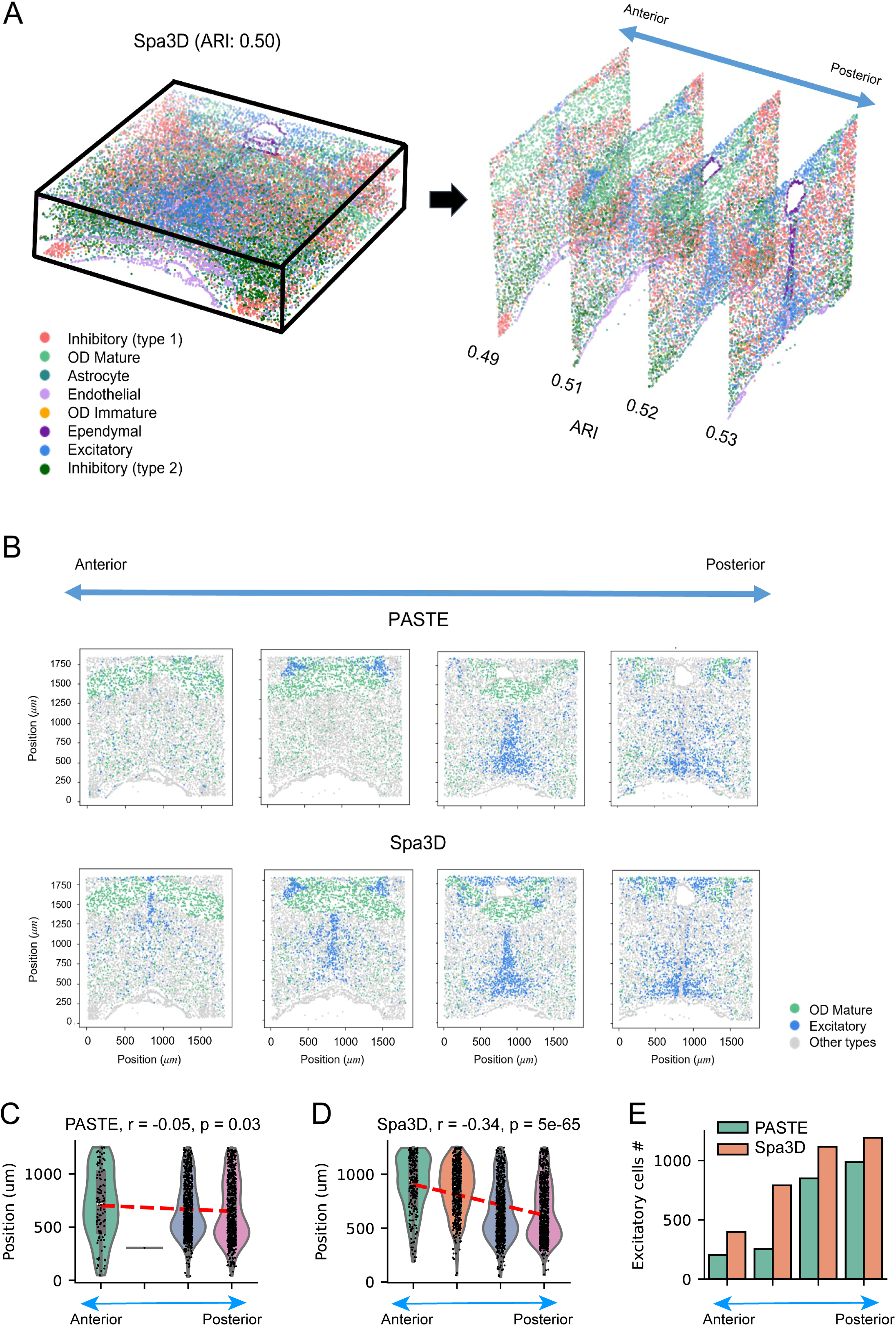
3D spatial trajectory in the mouse hypothalamus dataset based on the MERFISH platform. **(A)** The left panel displays the reconstruction of the 3D cellular organization by Spa3D. The distance between adjacent slices is 50 *µm*. For descriptive display purposes, we use different display scales for the z-axis and horizontal dimensions. The right panel presents the spatial trajectory from anterior to posterior. A planar layout of the Spa3D result is shown in Supplementary Figure 9. **(B)** Comparison of PASTE and Spa3D in presenting the spatial trajectory of OD mature and excitatory cells from anterior to posterior. **(C)** The total number of excitatory cells across the four slices in PASTE (green) and Spa3D (orange) representations. **(D-E)** The position distributions of excitatory cells in the lower half of the image across the four slices in PASTE **(D)** and Spa3D **(E)** representations. Only excitatory cells in the lower half were counted to display the trajectory because the cells in the upper half may belong to a separate group. There is no excitatory cell in the lower half of the second slice in PASTE representation. The red lines display the linear regression of vertical cell positions within the slices against the anatomical positions across slices from anterior to posterior.

To further examine the biological significance of Spa3D- and PASTE-defined spatial domains, we queried sdaSVGs for each annotated hypothalamus domain defined by Spa3D and PASTE with a known hypothalamus gene set defined by the Allen Mouse Brain Atlas^30^. As shown in **Supplementary Figure 11**, in most of these spatial domains, we observed that Spa3D-defined sdaSVGs showed higher overlap with known hypothalamus gene set than sdaSVGs defined by PASTE, by both edgeR (**Supplementary Figure 11A**) and Seurat algorithms (**Supplementary Figure 11B**). Moreover, several sdaSVGs defined by Spa3D but missed by PASTE (e.g., IGF1R^51,52^, ESR1^53–55^, PNOC^56^, and OXT^57,58^) have experimental evidence for hypothalamic function, validating Spa3D’s ability to detect histology-associated, domain-specific genes.

Second, one major advantage of Spa3D is the ability to uncover spatial trajectories in a 3D manner, which refers to the trajectory analysis that arranges cells based on their spatial patterns. This is achieved by aligning all cells across 2D SRT slices to reconstruct the intact 3D cellular organization (**Figure 5A, left panel**), which enables the visualization of 3D spatial trajectories of various cell types (**Figure 5A, right panel**). Conversely, stacking 2D slices in an arbitrary way without precise 3D alignment of all cells can result in inaccurate cell location determination, making it difficult to detect 3D spatial trajectories of different cell types. To compare the 3D spatial trajectory patterns revealed by Spa3D and PASTE in this hypothalamus dataset, we focused on two popular neuron types in the hypothalamic preoptic region: mature <u>o</u>ligo<u>d</u>endrocytes (OD mature) (green) and excitatory neurons (blue), to avoid too many cell types in one figure to mask the trajectory patterns (**Figure 5B**). We found that Spa3D and PASTE revealed similar patterns for MO cells after 3D spatial alignment and reconstruction (**green, Figure 5B**). However, for excitatory neurons, especially in the anterior hypothalamus, these two algorithms uncovered distinct patterns (**Figure 5B-E**): PASTE detected few excitatory neurons anteriorly and a punctuated posterior increase, whereas Spa3D identified more abundant and gradually increasing anterior excitatory neurons (**Figure 5C**). An independent EASI-FISH study^59^ confirmed the abundance and a gradually increasing gradient of excitatory neurons in hypothalamus, consistent with the findings by Spa3D. In addition, Spa3D better reveals the trajectory of spatial position change of excitatory neurons from anterior to posterior, as demonstrated in the linear regression results of vertical cell positions within slices against the anatomical positions across the slices from anterior to posterior (**Figure 5D-E**).

## DISCUSSION

Spa3D enables comprehensive 3D-based spatial segmentation analysis by incorporating spatial distances both across and within slices during 3D graph construction and GCN training. This framework naturally extends to 3D pseudotime inference and 3D cell-cell communication analysis (**Figures 2 and 3**). In addition, Spa3D supports 3D reconstruction from slices containing distinct cell types and heterogeneous gene expression patterns (**Figure 5**), demonstrating its applicability to complex biological tissues such as the brain and tumors. We further evaluated the performance of Spa3D on spatial transcriptomics (SRT) datasets generated by four major platforms, 10x Genomics Visium, ST, Stereo-seq, and MERFISH, which span a wide range of spatial resolutions. Spa3D can be readily applied to other emerging SRT technologies such as seqFISH+, STARmap, and Slice-seq, underscoring its versatility for diverse spatially resolved omics applications.

While Spa3D provides an effective framework for integrating multi-slice SRT data, its performance is partially influenced by the accuracy of inter-slice alignment. In this study, we either used the pre-aligned coordinates provided in the original publication (Figure 4) or applied the alignment methods implemented in PASTE (Figures 2, 3, and 5). Although techniques of 3D alignment across heterogeneous tissue sections have advanced, most existing approaches primarily rely on molecular features and often overlook complementary information contained in histological images. We envision that incorporating histological features alongside enhanced molecular signals will further improve 3D alignment fidelity. The improved version of Spa3D in our follow-up study will leverage cellular components, morphological features, and tissue architecture to guide alignment, ensuring cross-slice consistency and preserving local tissue integrity. This improvement will enable more accurate 3D reconstructions and provide deeper biological insights.

Beyond spatial transcriptomics, Spa3D offers a foundation for extending 3D reconstruction to other spatial omics modalities, including spatial epigenomics (e.g., spatial ATAC-seq and spatial-CUT&Tag) and spatial proteomics. Such extensions would enable exploration of epigenomic regulation and protein interaction networks within true 3D tissue contexts. By restoring tissue morphology and embedding spatial coordinates into multi-omics data, Spa3D establishes a general framework for advancing our understanding of cellular function, tissue development, and disease progression in three-dimensional space.

## METHODS

### Spatial pattern enhancement (SPE)

We have used two approaches below to enhance the spatial pattern. First, we have transformed the gene expression from the real-number domain to the complex-number domain. We first calculated the analytic expression from the real-number expression:

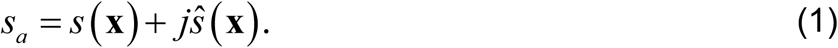

Here, the *s* is the gene expression at spatial point **x**, *ŝ* is the Hilbert transform of *s*. *s_a_* is the analytic signal. We then calculated the envelop *s_e_* of the gene expression from the analytic signal:

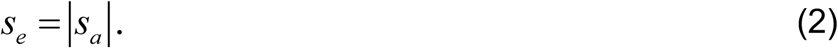

Considering the gene expression is the oscillating signal, the envelope of the signal consists of its local extreme values. It generalizes the local gene expression values into instantaneous gene expression.

This SPE approach works effectively for the DLPFC dataset (**Figure 2**), in which the SRT samples are sufficient, and the spatial structures are well organized. However, the human embryonic heart dataset (**Figure 3**) contains fewer SRT samples, and the mouse hypothalamus dataset (**Figure 4**) provide single-cell resolution. To control the degree of enhancement under such varying conditions, we developed a second SPE approach that applies the anti-leakage Fourier transform (ALFT) to extract the dominant components of spatial patterns. The ALFT was proposed by Xu et al. (2005, 2010)^60,61^ for seismic data regularization. It was later used by Tang and McMechan (2017)^62^ to extract the normal direction of reflection image in local windows.

In a local spatial window, we use the forward Fourier transform to project the real-domain signal, i.e., the gene expression *s*(*x*), into the wavenumber domain:

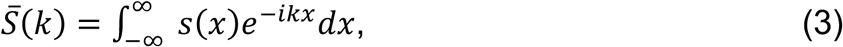

where *S̅*(*k*) denotes the complex Fourier coefficient of the gene expression *s*(*x*) at wavenumber *k*. If a signal has spatial structure, its energy tends to concentrates at specific wavenumbers. In contrast, spatially unstructured signals (e.g. random noise or point sources) exhibit energy dispersed across a wide range of wavenumbers. This property underlies the ALFT algorithm’s ability to isolate meaningful spatial patterns.

ALFT works by iteratively extracting the highest-energy component in the Fourier domain until the residual signals in the real domain fall below a set threshold:

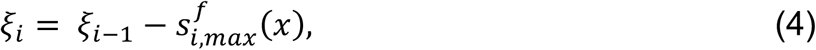

where *ξ_i_* and *ξ_i_*_− 1_ are the residual signals at the current iterations *i* and *i* − 1, respectively. The initial residual is defined as:

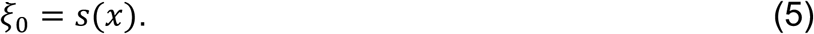

Here, 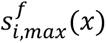 is the inverse Fourier transform of the dominant frequency component 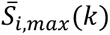:

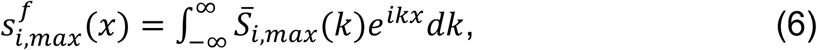

where 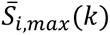 is the complex Fourier coefficient at the wavenumber *k*with the largest magnitude at iteration *i*, corresponding to the dominant spatial frequency component extracted at that step. By successively extracting these high-energy components and updating the residual, ALFT progressively emphasizes coherent spatial structures while suppressing unstructured background noise.

Equations 4 to 6 perform well when spatial patterns are strong, and wavenumber values are sparse. However, gene expression patterns are typically weaker and less sparse than seismic signals, making the iterative inverse Fourier transform computationally expensive. To improve efficiency, we perform the iteration in the wavenumber domain:

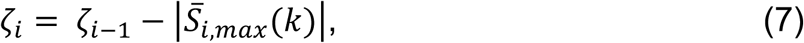

where *ζ_i_* is the residual energy in the wavenumber domain at iteration *i*, and

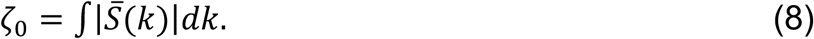

This approach avoids repeated inverse transforms while maintaining the same iterative filtering principle. We could optionally square the 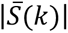 terms in equations 7 to 8, but this is not necessary since the iteration already achieves effective filtering of dominant wavenumbers.

After identifying all effective wavenumber points, we perform a single inverse Fourier transform to map them back to the spatial domain, avoiding repeated transforms during each iteration. This is an efficient way for the gene expression as the value in the wavenumber domain is often not sparse. If the energy threshold *ζ_threshold_* is small (e.g., 5% of the total energy), the iteration can alternatively accumulate the lowest energy components:

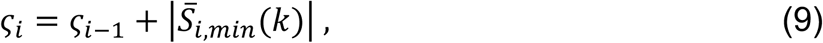

with

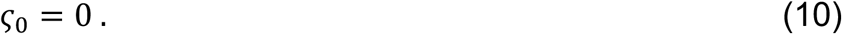

This alternative follows the same principle as *ζ_i_*, but in reverse, and provides flexibility for controlling the retained energy. One can also rank all wavenumber magnitudes before iteration and select the top or bottom entries at each step.

Overall, the ALFT-based spatial pattern enhancement iteratively identifies and extracts the dominant spatial frequency components in the wavenumber domain. Each wavenumber *k* corresponds to a particular spatial variation in the real domain, and by combining these components, the method reconstructs the prominent spatial patterns. By suppressing lower-energy components, which typically represent noise, the process enhances the primary spatial structures while simultaneously performing denoising.

To empirically validate the assumption that spatial signals concentrate energy at specific wavenumbers, we computed the fraction of Fourier power in the top 5% of wavenumbers (Top5PercentEnergy) across all genes in four spatial omics datasets from the heart. We observed that a substantial portion of genes (47.09% to 49.70% across datasets) have Top5PercentEnergy exceeding 0.9, with maximum values reaching 0.999, while many other genes exhibit moderately concentrated energy. These results indicate that nearly half of the genes carries highly concentrated spatial signals, whereas others are more evenly distributed, consistent with the expected diversity of spatial structures across genes. Overall, this analysis provides quantitative support for the principles underlying ALFT- and Hilbert transform-based spatial pattern enhancement in our pipeline.

### 3D reconstruction based on graph convolutional network (GCN)

For 3D reconstruction, as each slice may have inter-slice registration errors, we first applied pairwise alignment implemented in PASTE^24^ to the multiple 2D SRT slices before assembling the 3D dataset. The 3D data can be used for both 3D reconstruction and pseudotime inference analysis (e.g., Figures 2A and 2C). We then utilized the full 3D spatial information, including the depth z coordinate, to calculate the adjacency matrix *A* for graph construction. When 3D histology images are involved in calculating the Euclidean distance, Spa3D provides an option to apply a penalty parameter σ to the depth z coordinate to correct for anisotropic resolution across dimensions:

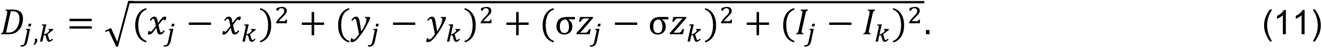

where the *j* and *k* denote two spots, (*x, y, z*) is the 3D coordinate of a spot, and the *I* is a distance measured by the RGB value of the histology image in a local window. In typical spatial transcriptomics data, the z-axis (across slices) is sampled more coarsely than the x-y plane, leading to lower effective resolution along depth. To compensate, we scale the z values using σ = (*l_c_* + *d*)/*l_c_*, where *l_c_* is the center-to-center distance between adjacent spots within a single slice (100 µm) and *d* is the spot diameter (55 µm) in the 10x Genomics Visium platform, giving σ = 1.55. This scaling increases the contribution of z-axis differences to the Euclidean distance, correcting for lower axial resolution. If isotropic resolution is assumed (σ = 1), Equation 11 reduces to the standard 3D Euclidean distance:

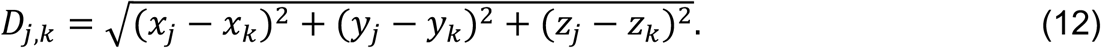

The *D_j,k_* will be used to construct the adjacent matrix A which consists of *A_ij_*:

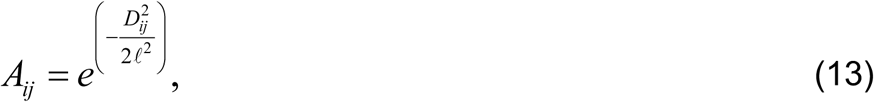

where *A_ij_* is the edge weight between nodes *i* and *j*, and £ is a parameter that determines the scale of the kernel, affecting how quickly the values decay with increasing distance. We used Louvain for the initialization of clustering analysis and used GCN to iteratively update the 3D reconstruction by optimizing the lost function. The graph convolution is

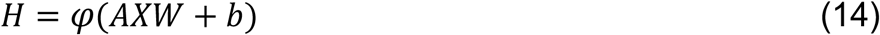

where *A* is the adjacent matrix calculated using full 3D coordinates (referring to equations 11 and 12), *X* is the matrix of the input features, *W* is the weight matrix of the layer, *b* is the bias vector, φ(·) denotes the nonlinear activation, and *H* is the output. The loss function uses Kullback-Leibler (KL) divergence:

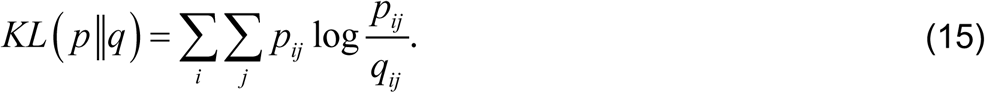

Here, *q* is the predicted soft assignment distribution, representing how likely each embedded data point is to belong to a given cluster. Mathematically, *q* is calculated using a Student’s *t*-distribution with one degree of freedom (also known as the Cauchy distribution) as a kernel to measure the similarity between the *i*-th embedded data point **z***_i_* and the *j*-th cluster center *μ_j_*:

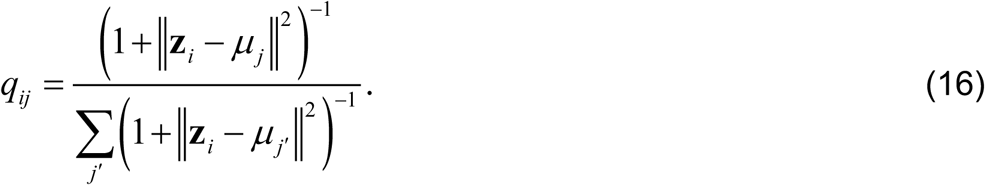

In equation 15, *p* is the target distribution, used to guide the optimization of the clustering by serving as a reference or “target” for updating the model’s parameters. The target distribution is calculated in a way that emphasizes data points with high confidence in their cluster assignment and minimizes the entropy of the distribution. *p* can be calculated from *q*:

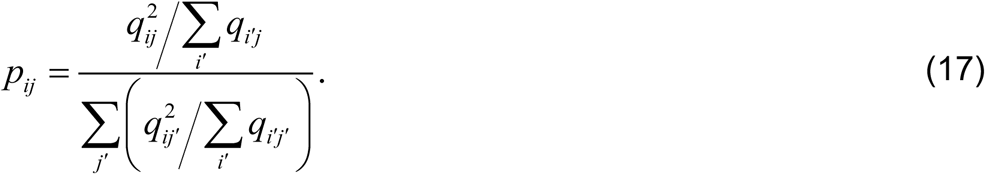

The Kullback-Leibler divergence between *q* and *p* is often minimized during training.

Equation 14 is a standard GCN layer formulation^63^, and equations 15 to 16 represent the routine process for implementing unsupervised deep embedding for clustering analysis^64^. The approach is used by Hu et al.^8^. The main difference in our approach is that we use 3D coordinates to calculate the adjacency matrix for the graph, which is then used in the subsequent calculations.

### DLPFC dataset

We selected the first approach, instantaneous amplitude, to enhance the spatial pattern due to the large sample size. To be consistent with the PASTE method^24^, we used the original data provided from the PASTE GitHub page to generate the PASTE-defined “center slice” in **Figure 2B**.

### Human heart dataset

The annotation was obtained from Asp et al. (2019) ^37^ by merging similar, smaller clusters into five larger ones. As the PASTE paper does not provide the “center slice” result for this example, we used the PASTE scripts from its GitHub page to generate the integrated “center slice” displayed in **Figure 3B**. We applied a SpaceFlow-based DPT^65^ for the pseudotime inference analysis in **Figure 3E**, using 3D spatial coordinates and time information.

### *Drosophila* embryos dataset

The annotation was obtained from the original paper^50^. We selected the slices 9-11 in E14-16 stage for analysis. The dataset contains both raw pixels and pre-aligned positions. For Spa3D analysis, we used the raw pixels to do spatial enhancement and the pre-aligned positions to do domain segmentation. For PASTE analysis, we used PASTE-aligned positions to do domain segmentation.

### Hypothalamic preoptic region dataset

Because the dataset provided by the MERFISH technology is at the single-cell level, we used the second approach, revised ALFT in local windows, to enhance the local spatial pattern.

### Downstream analysis of Spa3D and PASTE results

We calculated pseudotime by running the diffusion pseudotime (DPT) method^21^ using the Spa3D and PASTE embedding, respectively. For cell-cell communication analysis, we used two distinct frameworks, LIANA^33^ and SpaTalk^14^, for independent analysis. To define domain-specific spatially variable genes (SVGs), we performed differential expression (DE) analysis using spots from a target domain versus all other domains. To achieve a robust list of SVGs, we applied edgeR^28^ and Seurat^29^ packages, both of which are state-of-art DE analysis methods.

## Code availability

The Spa3D package is available at https://github.com/Lin-Xu-lab/Spa3D.git.

## Acknowledgments

The resources of the high-performance computing environment from the Quantitative Biomedical Research Center (QBRC) and BioHPC at UT Southwestern Medical Center, as well as the Texas Advanced Computing Center (TACC) at The University of Texas at Austin, are gratefully acknowledged. We also thank Ms. Jessie Norris for proofreading this manuscript.

## Contributions

CT, YZ, XX, and LX conceived and designed the study. CT, YZ, and XX developed the Spa3D algorithm, generated the scripts and GitHub page, and performed the data analysis. LD and LY were involved in the data analysis. LX and GX acquired the funding. CT, YZ, XX, QL, GX, and LX wrote and revised the manuscript. All authors have read, revised, and approved the final manuscript.

## Ethics declarations

All the authors declare that they have no competing interests.

## Data availability

The datasets used in this study are publicly available, and the original publications for each dataset are cited in the Methods section. The source code for Spa3D can be downloaded from GitHub (https://github.com/Lin-Xu-lab/Spa3D.git)

