## Supplementary Figure for "3D reconstruction of spatial transcriptomics with spatial pattern enhanced graph convolutional neural network"

**Supplementary Figure 1**

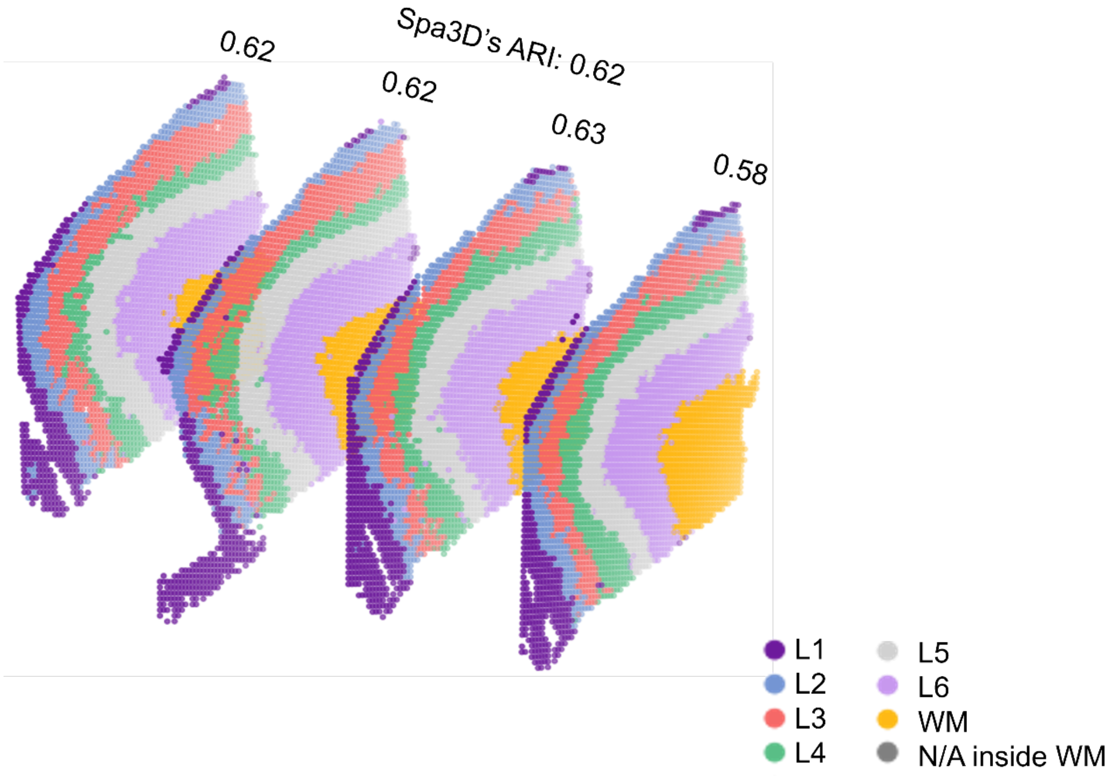


**Supp. Figure 1**: All of the Spa3D-reconstructed individual slices (ARI from 0.58 to 0.64) outperformed this PASTE-defined “center slice” in clustering accuracy, not just the one showcased in Figure 2B.

**Supplementary Figure 2**

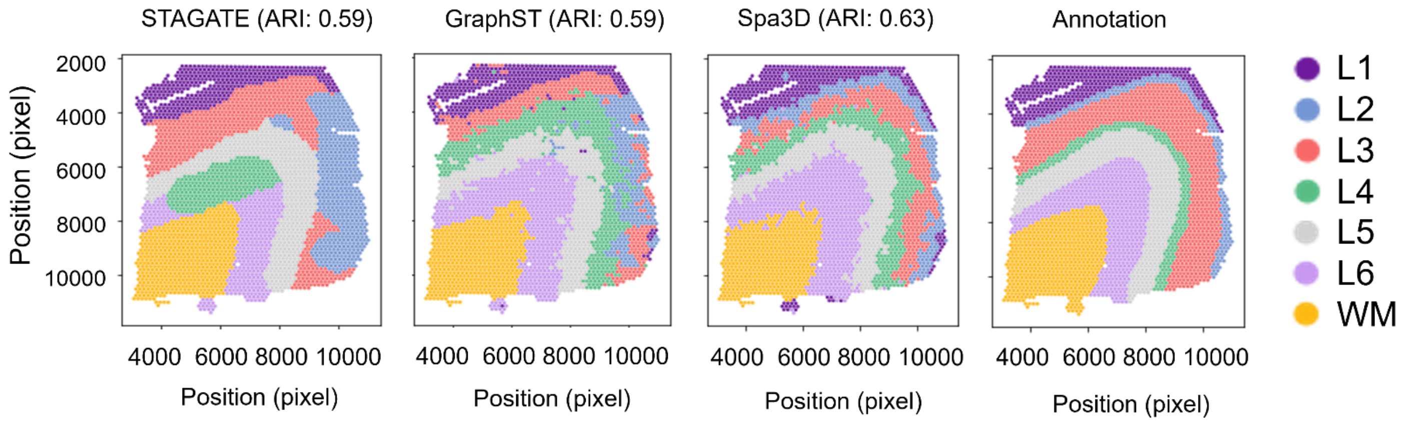


**Supp. Figure 2:** Comparison between STAGATE, GraphST, and Spa3D results. This comparison is a supplement to Figure 2B using the human dorsolateral prefrontal cortex dataset. The green cluster in STAGATE is an undefined cluster, as it does not spatially correspond to any cluster in the annotation.

**Supplementary Figure 3**

A

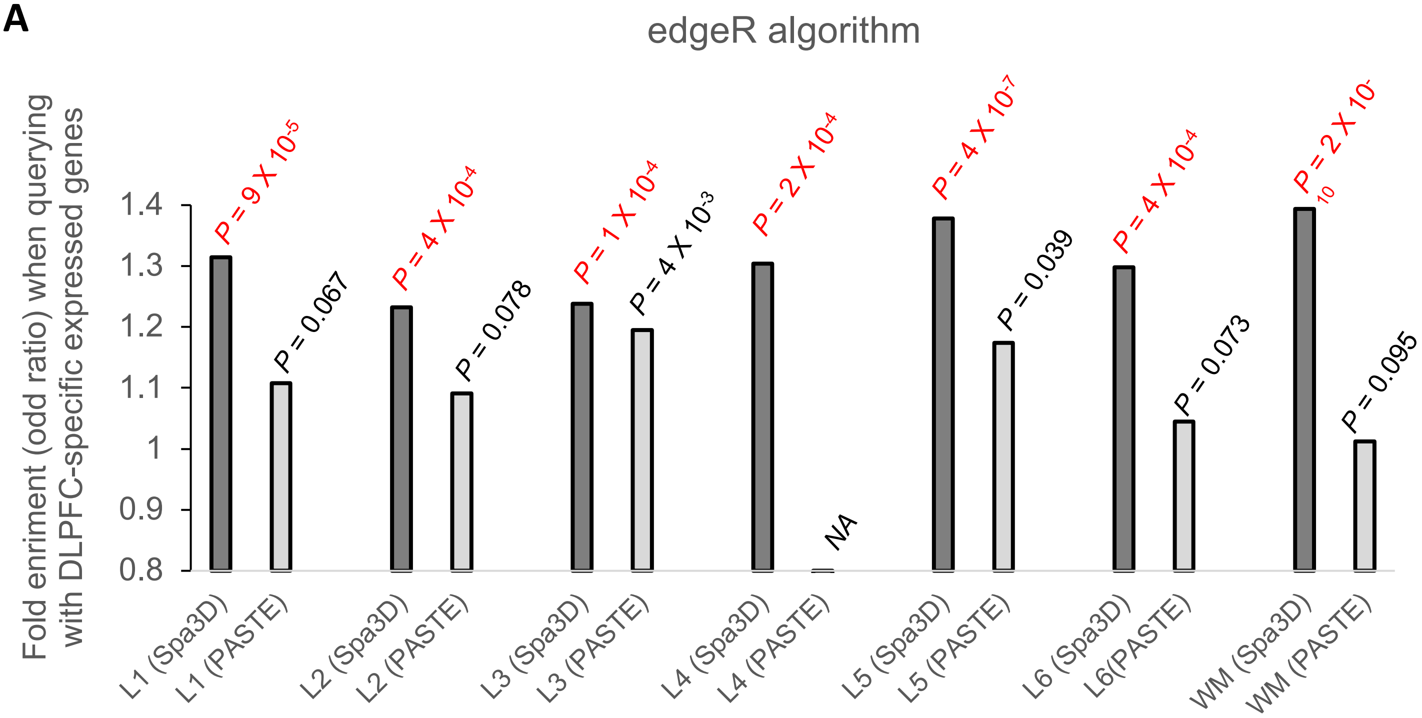


B

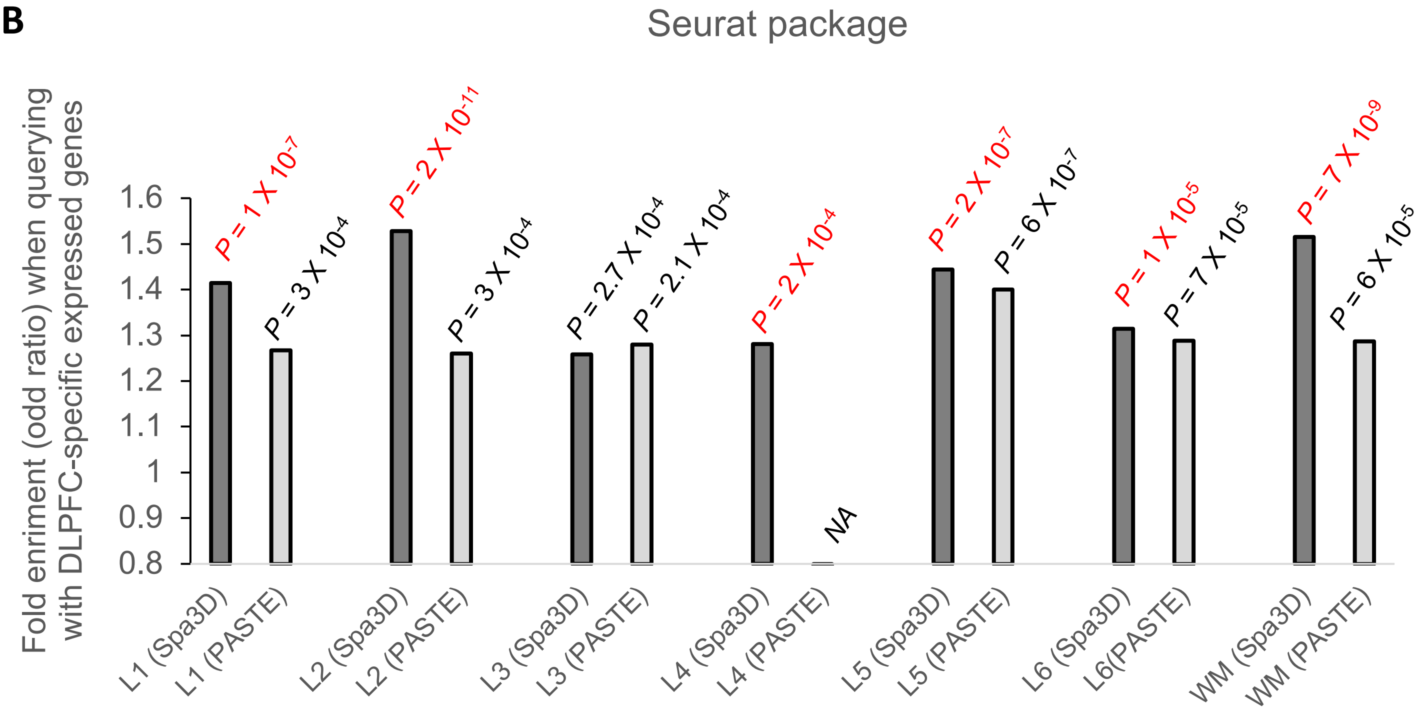


**Supp. Figure 3**: Overlap between spatially variable genes of each individual DLPFC layer (sdaSVGs) with known DLPFC genes by edgeR method (**A**) and Seurat package (**B**). P values of enrichment analysis were shown in the figures. Whenever Spa3D-defined sdaSVGs display more significant enrichment with known DLPFC genes (that is, lower P value) than PASTE-defined sdaSVGs, we used red color to label P value for the Spa3D bar. It is worth noting that PASTE could not detect L4 layer in DLPFC samples but Spa3D can (**Figure 2B**), and therefore a “NA” label for PASTE was added to represent this discrepancy.

**Supplementary Figure 4**

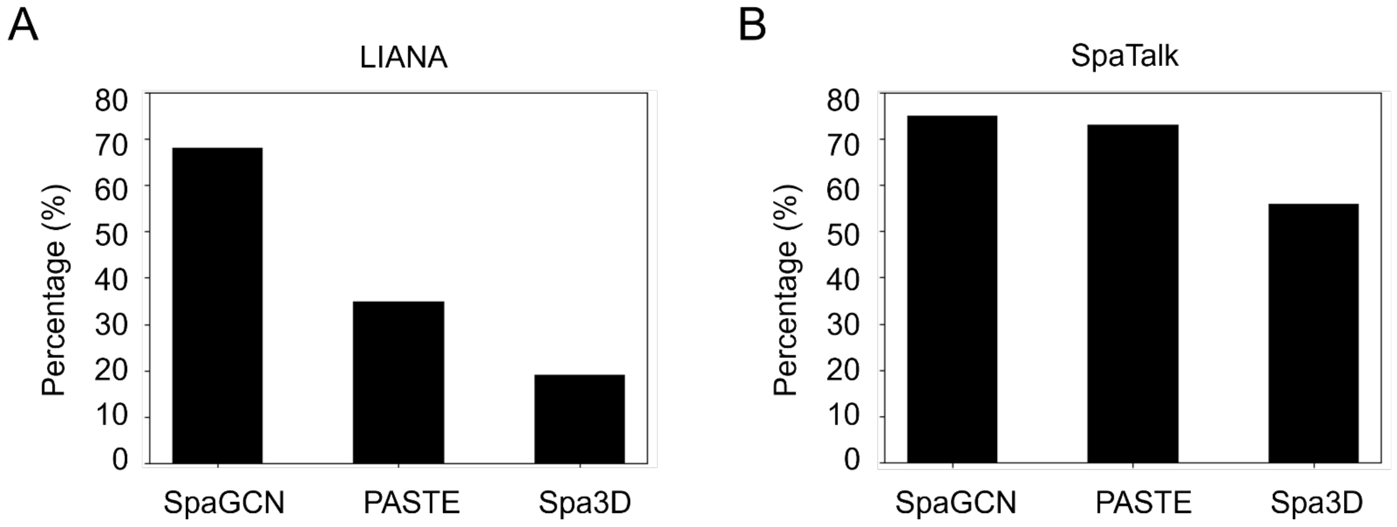


**Supp. Figure 4:** Comparison of the false positive rate (FPR) for ligand-receptor pairs (FPLRPs) across the results from SpaGCN, PASTE, and Spa3D, analyzed using LIANA (**A**) and SpaTalk (**B**) on the DFPFC dataset. For both LIANA and SpaTalk analyses, Spa3D exhibits lower FPRs compared to SpaGCN and PASTE.

**Supplementary Figure 5**

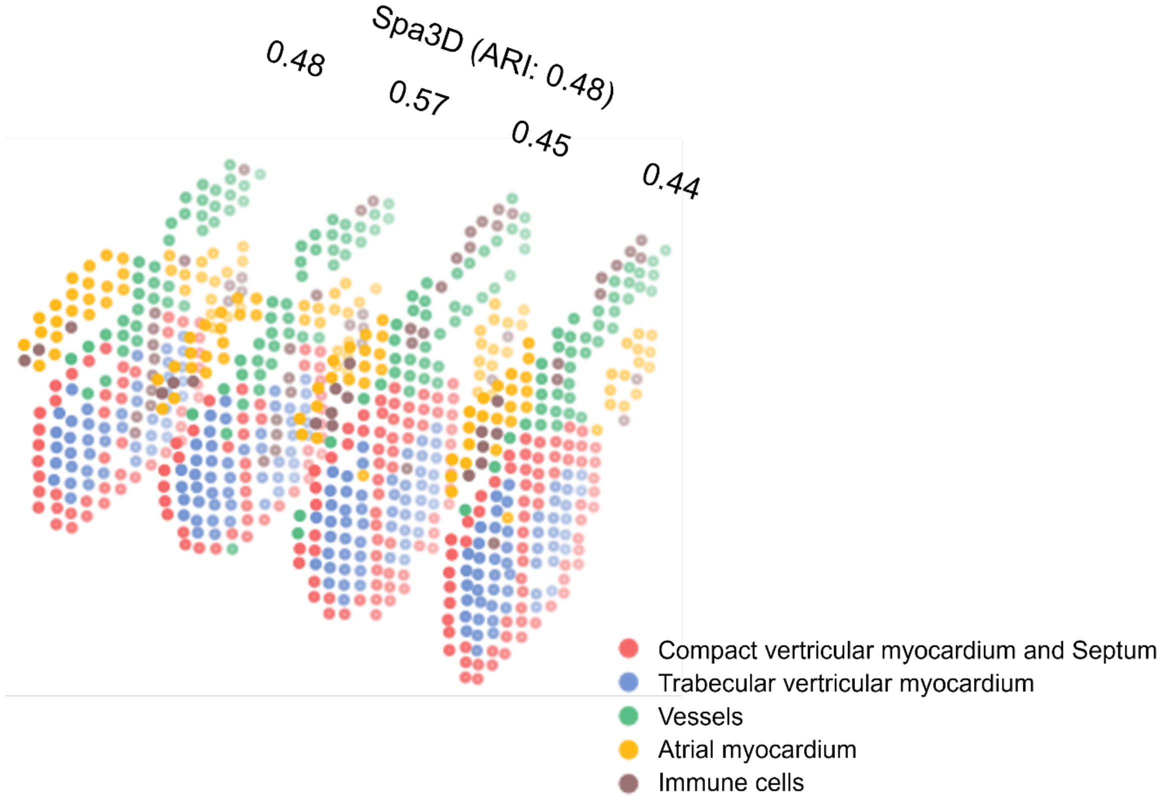


**Supp. Figure 5**: All individual slices reconstructed by Spa3D (ARI from 0.44 to 0.57) achieved higher ARI scores than PASTE-defined center slice, not just the one showcased in Figure 3B.

### Supplementary Figure 6

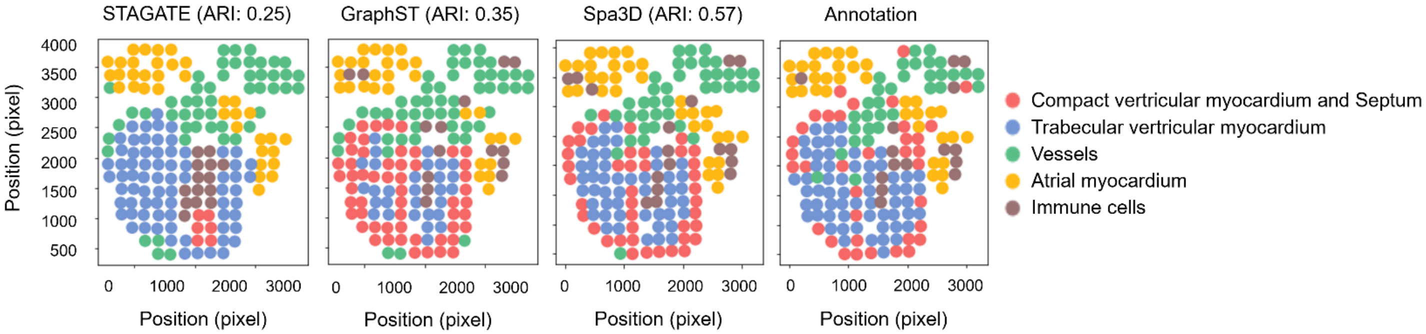


**Supp. Figure 6:** Comparison between STAGATE, GraphST, and Spa3D results. This comparison is a supplement to Figure 3B using the human embryonic heart dataset.

**Supplementary Figure 7**

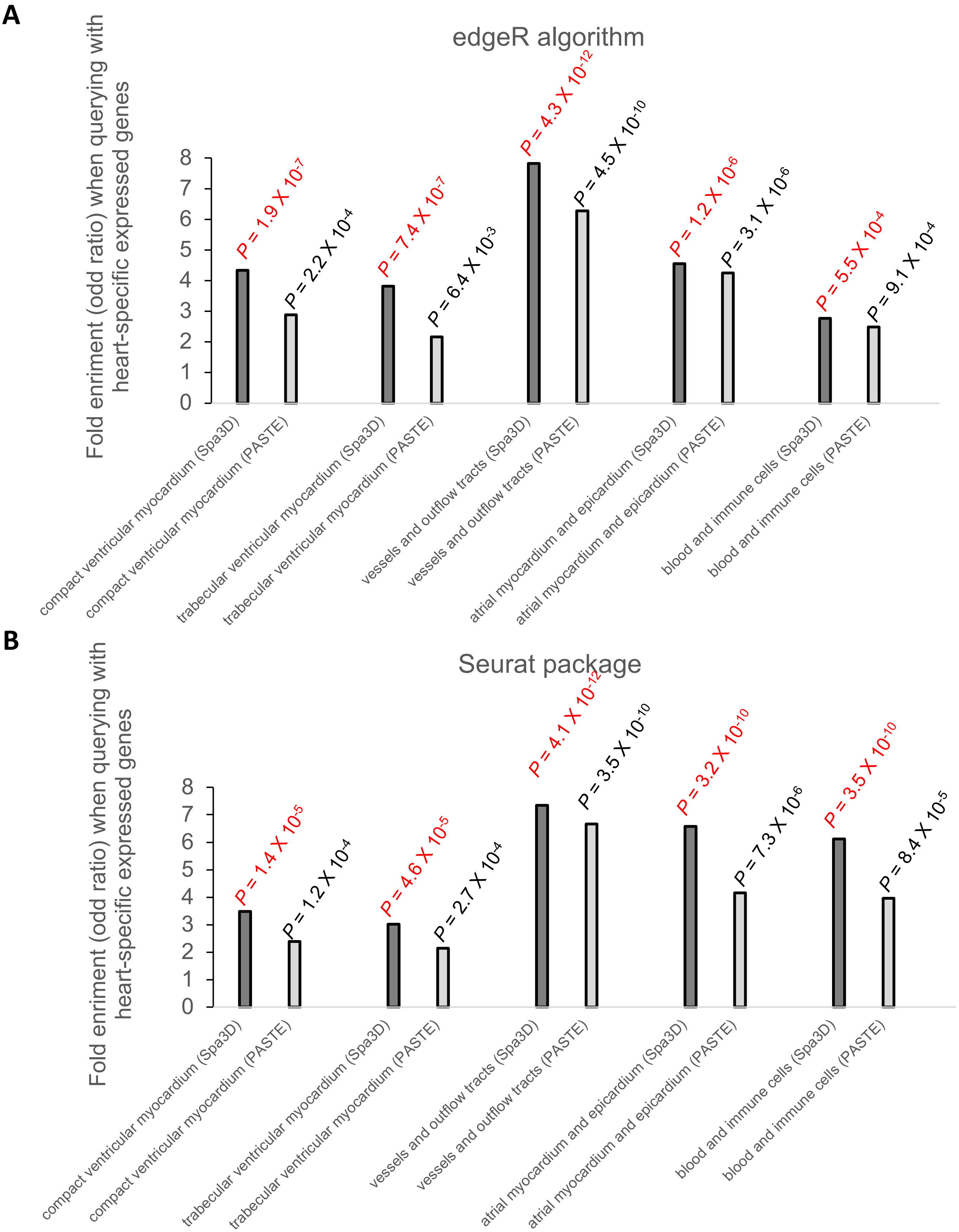


**Supp. Figure 7**: Overlap between sdaSVGs of each individual heart domain with known heart-specific tissue type genes by edgeR method (**A**) and Seurat package (**B**). Whenever Spa3D-defined sdaSVGs display more significant enrichment with known genes, we used red color to label P value.

**Supplementary Figure 8**

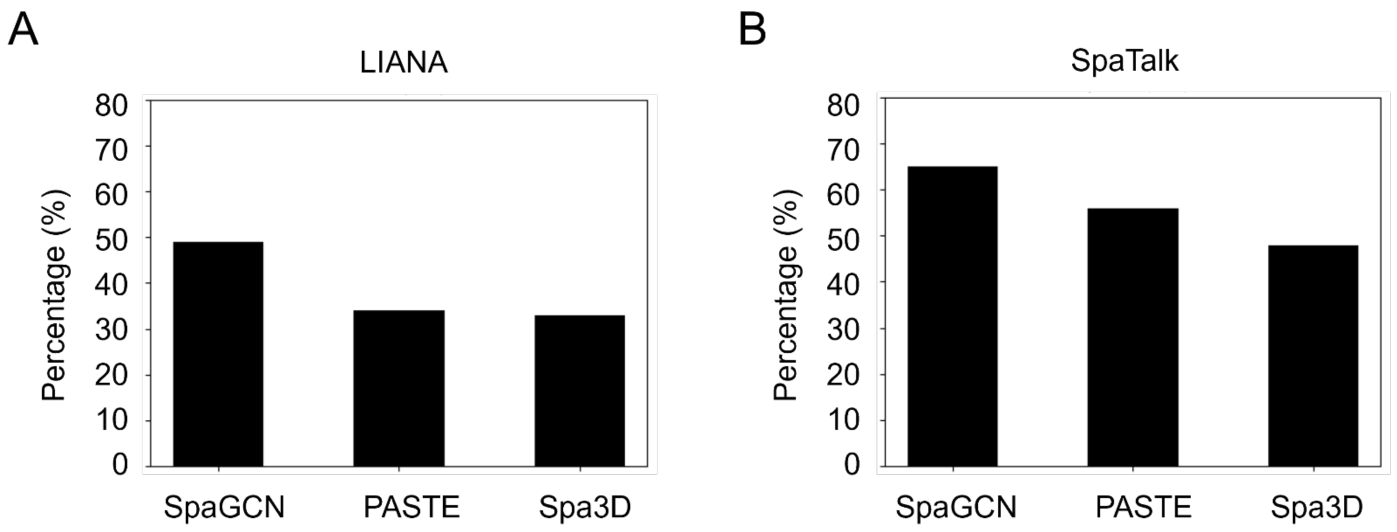


**Supp. Figure 8:** Comparison of the false positive rate (FPR) for ligand-receptor pairs (FPLRPs) across the results from SpaGCN, PASTE, and Spa3D, analyzed using LIANA (**A**) and SpaTalk (**B**) on the human embryonic heart dataset. For both LIANA and SpaTalk analyses, Spa3D exhibits lower FPRs compared to PASTE and SpaGCN.

**Supplementary Figure 9**

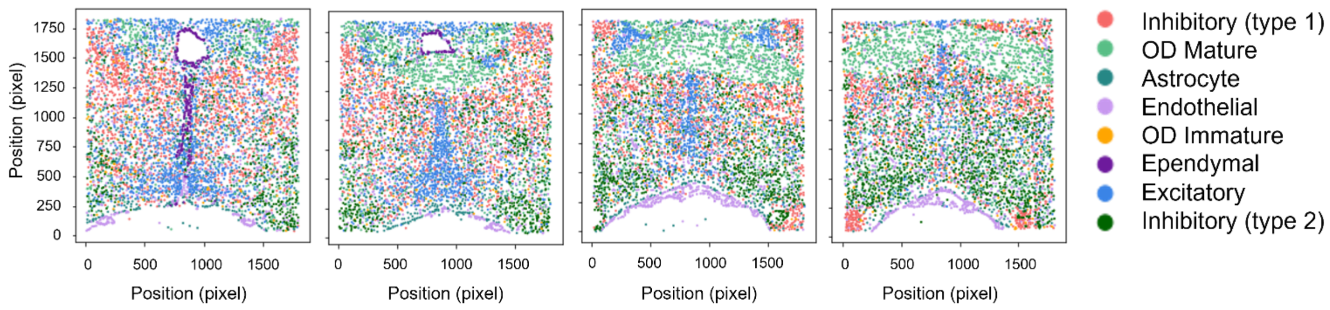


**Supp. Figure 9**: The planar layout of the Spa3D result. This figure is a supplement to Figure 4A.

**Supplementary Figure 10**

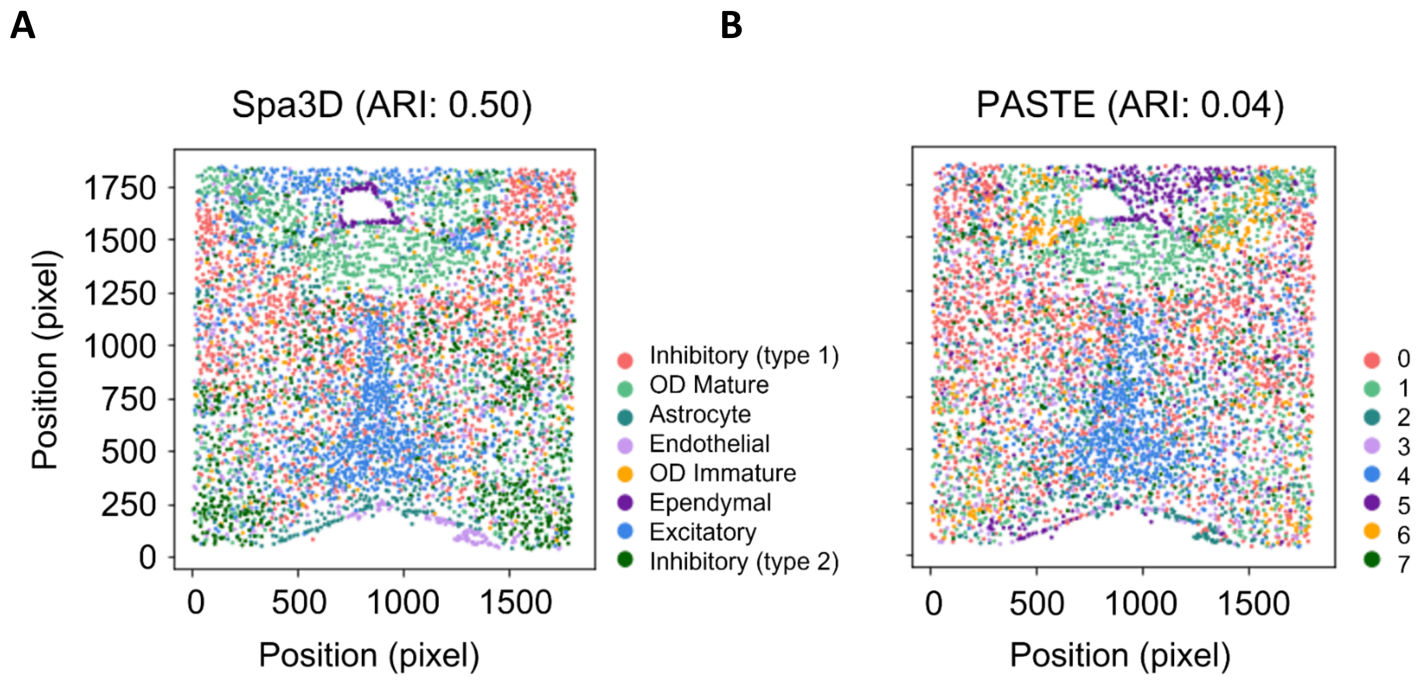


**Supp. Figure 10**: Comparison of PASTE-integrated "center slice" and the corresponding slice from Spa3D. (**A**) The corresponding slice from Spa3D. The slice ID is 2. (**B**) PASTE-integrated "center slice"**.**

**Supplementary Figure 11**

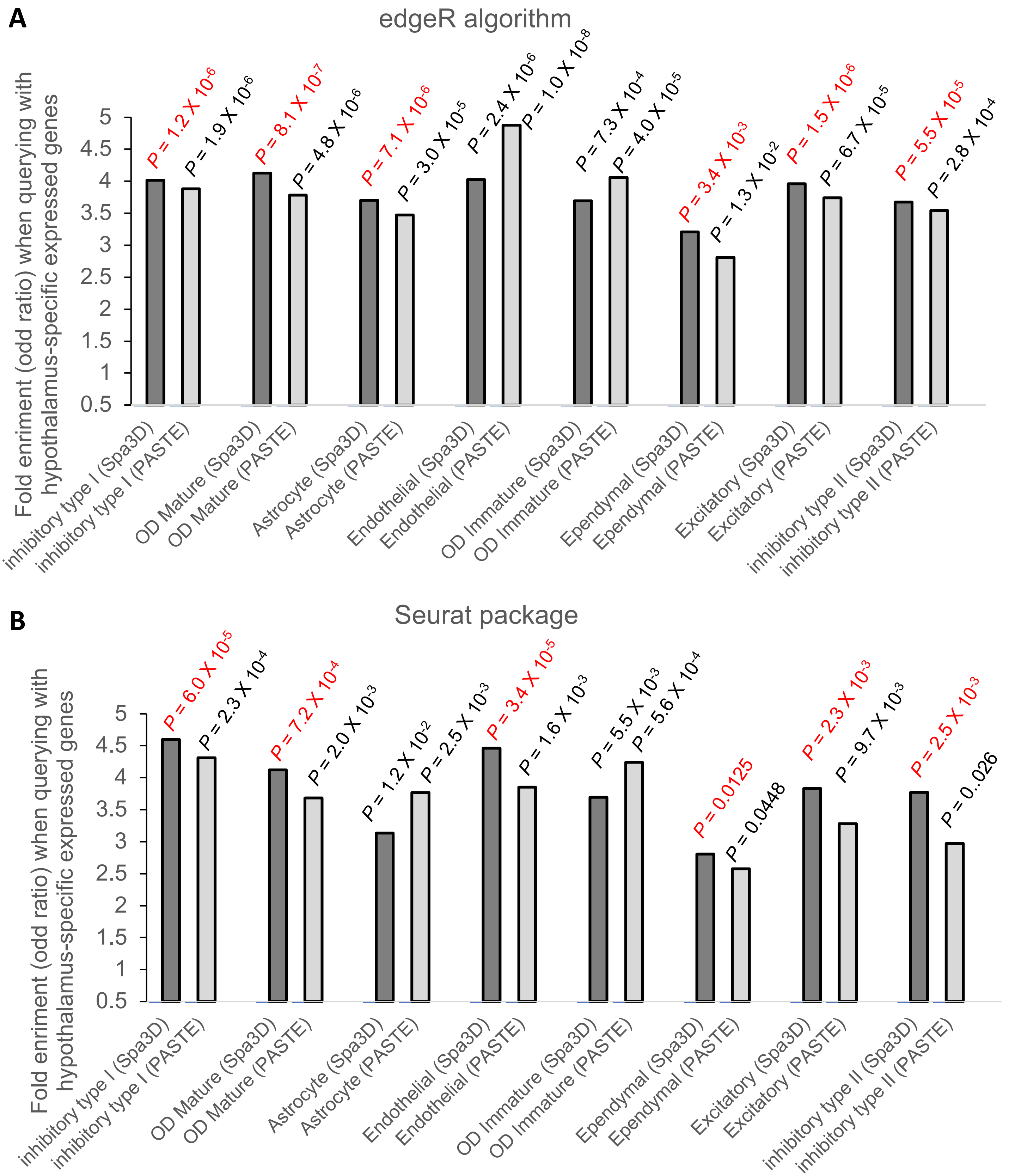


**Supp. Figure 11**: Overlap between sdaSVGs of each individual hypothalamus domain with known hypothalamus genes by edgeR method (**A**) and Seurat package (**B**). Whenever Spa3D-defined sdaSVGs display more significant enrichment with known hypothalamus genes (that is, lower P value) than PASTE-defined sdaSVGs, we used red color to label P value for the Spa3D bar.
